# A Conceptual Framework for Host-Associated Microbiomes of Hybrid Organisms

**DOI:** 10.1101/2023.05.01.538925

**Authors:** Benjamin T. Camper, Zachary Laughlin, Daniel Malagon, Robert Denton, Sharon Bewick

## Abstract

Hybridization between organisms from evolutionarily distinct lineages can have profound consequences on organism ecology, with cascading effects on fitness and evolution. Most studies of hybrid organisms have focused on organismal traits, for example various aspects of morphology and physiology. However, with the recent emergence of holobiont theory, there has been growing interest in understanding how hybridization impacts and is impacted by host-associated microbiomes. Better understanding of the interplay between host hybridization and host-associated microbiomes has the potential to provide insight into both the roles of host-associated microbiomes as dictators of host performance as well as the fundamental rules governing host-associated microbiome assembly. Unfortunately, there is a current lack of frameworks for understanding the structure of host-associated microbiomes of hybrid organisms. In this paper, we develop four conceptual models describing possible relationships between the host-associated microbiomes of hybrids and their progenitor or ‘parent’ taxa. We then integrate these models into a quantitative ‘4H index’ and present a new R package for calculation, visualization, and analysis of this index. Finally, we demonstrate how the 4H index can be used to compare hybrid microbiomes across disparate plant and animal systems.

## 1. Introduction

Hybridization is increasingly recognized as an important component of ecological and evolutionary processes. Consequences of hybridization span the fitness spectrum, ranging from infertility and death^1, 2^ on one end to innovation and adaptation on the other.^3–6^ Ultimately, it is the balance of the positive and negative outcomes of hybridization that dictates the role that hybridization plays in the success or failure of different genetic lineages.^5, 7–9^ If, for example, hybridization produces sterile offspring, then it can drive the emergence of genetic sinks and evolutionary dead ends^10^ and thus serve as a ‘brake’ for evolution. Alternately, if hybridization facilitates ecological release and/or sexual isolation (either directly through mating barriers or indirectly through altered temporal or spatial proximity), then it can promote lineage diversification and thus serve as a ‘motor’ for evolution.^11^

Most early research on hybrid organisms focused on understanding how hybridization impacts host fitness though effects on host traits, for example, fecundity^12–16^, physiology^17–21^, morphology^22–25^ and behavior.^26, 27^ Recently, however, there has been growing recognition that macroorganisms are not autonomous units. Rather they are collectives or ‘holobionts’ compromised of both a host and all of its host-associated (HA) microbes.^28–31^ Thus, just as it is important to understand how hybridization impacts the traits of the host, it is equally important to understand how hybridization impacts the traits of the holobiont, including characteristics of the host-associated microbiome.^32^ Indeed, the eco-evolutionary basis for holobionts has led to entirely new branches of research in areas as diverse as human health^33, 34^, conservation^35, 36^ and biotechnology,^37, 38^ and it is currently poised to do so within the field of hybridization research as well.

The importance of the holobiont stems from the many host traits and processes that are either partially or fully dependent on host-associated microbes.^39–42^ As an example, gut microbiomes are strong regulators of host metabolic phenotype.^43–45^ This, in turn, impacts host energy balance,^46–48^ including both energy intake and use and expenditure. Gut microbiomes can also be important determinants of dietary niche,^49–53^ either by provisioning hosts with key nutrients^54–56^ or by detoxifying defensive compounds found in host food sources.^57^ Beyond diet and metabolism, HA microbiomes influence a range of other host traits as well.^58–68^ Healthy gut^69, 70^, skin^69, 71, 72^, and vaginal microbiomes^73^, for example, provide pathogen resistance across a broad spectrum of animal species.^74–76^ Indeed, amphibian skin microbiomes have been extensively studied as a means of defending hosts from devastating fungal pathogen (*Batrachochytrium dendrobatidis* and *B. salamandrivorans*) epidemics.^77–80^ In humans, disruptions to healthy HA microbiomes also underly a range of noninfectious diseases^81, 82^ such as rheumatoid arthritis^83, 84^ and irritable bowel syndrome.^85, 86^ Ultimately, the cascading effects of HA microbiomes on host traits and processes – ranging from host energy balance and dietary niche through to disease risk and immune dysfunction – have strong consequences on host ecological success^87^ and, by extension, host evolution.^88–90^

Although there has been substantial literature documenting both coevolutionary^91–95^ processes and codiversification patterns^94, 96–98^ between hosts and their HA microbiomes^99–108^, the study of how HA microbiomes respond when divergent host lineages reunite, or admix, through hybridization is relatively new.^109^ One of the earliest investigations into hybrid microbiomes was in *Nasonia* wasps.^1^ In this system, up to 90% lethality is observed in *F*_2_ males of *N. vitripennis/N. giraulti* crosses. However, rearing wasps under germ-free conditions results in near complete rescue of the same *F*_2_ males. This suggests a microbial basis to hybrid lethality. Interestingly, the 10% of hybrid *N. vitripennis/N. giraulti* males that survive under natural conditions exhibit highly transgressive microbial phenotypes. This includes both the appearance of novel microbial taxa in hybrid microbiomes as well as shifts in the abundances of microbial taxa that are shared among parents and hybrids.

More recent studies on hybrid vertebrates paint a similar picture. For example, hybrid house mice (*Mus musculus musculus* and *Mus M. domesticus*) in central Europe^110^ exhibit widespread transgressive microbiomes. Further, like the *Nasonia* wasp system, there is evidence that the altered microbial phenotypes of hybrid individuals at least partially explain their poor fitness outcomes.^111–116^ In particular, there is an interaction between inflammation, immune gene expression and the gut microbiome that appears to cause hybrid mice to exhibit defects in immunoregulation. This may be one reason why hybrid individuals are restricted to a narrow tension zone where the two parent subspecies co-occur.^117, 118^ A range of additional studies, including hybridization of sika deer (*Cervus nippon*) and elk (*Cervus elaphus*)^119^, lake whitefish lineages (*Coregonus clupeaformis*)^120^, blunt snout bream (*Megalobrama amblycephala*) and topmouth culter (*Culter alburnus*),^121^ and desert (*Neotoma lepida*) and Bryant’s (*Neotoma bryanti*) woodrats^122^ have reiterated the finding that hybrid animals often exhibit altered microbiomes relative to their progenitors (i.e., ‘parent’ lineages or ‘parent’ taxa). Indeed, even beyond the animal kingdom, hybrid macroorganisms are commonly associated with perturbations to the HA microbiome.^123–125^

As suggested above, the study of hybridization and its impact on HA microbiomes is important for understanding host fitness and evolution.^126, 127^ But even beyond questions of host success, hybrid systems are of interest because they facilitate an understanding of genotype-phenotype interactions.^128, 129^ Many hybrid zones^21, 26, 130^, especially systems where *F*_2_ individuals readily admix with their progenitors, provide variable genetic combinations^131, 132^ and degrees of heterozygosity across hybrid individuals. Consequently, these systems serve as natural laboratories for understanding how host genetics and environmental characteristics influence host traits. For example, in a study investigating how host genetics and the environment impact HA microbiomes across a *Neotoma* woodrat hybrid zone, Nielson *et al.* (2023)^122^ demonstrated that HA microbial composition was predominately driven by host genetics (genotypic classes), while HA microbial richness was predominately driven by the environment (core diet + vegetation communities). Applying similar approaches to other hybrid systems may be a fruitful avenue for disentangling the long-standing nature versus nurture paradigm as it applies to HA microbiomes and HA microbiome assembly.

Despite the increasing recognition that HA microbiomes are an important facet of hybridization and that hybrid organisms are valuable systems for understanding HA microbiome structure and function, there is a lack of frameworks for describing and comparing hybrid HA microbiomes across the tree of life. In this paper, we develop four conceptual models delineating potential relationships between hybrid microbiomes and the microbiomes of their progenitors. We discuss the underlying implications of each model, how each model might arise based on fundamental host mechanisms, and how each model could impact host fitness. We then integrate these four models into a quantitative ‘4H index’ that can be used to assess the relative importance of each model across widely disparate hybrid systems. Finally, we introduce an R package, `HybridMicrobiomes`, containing a series of functions that allow researchers to apply the 4H index to their own hybrid microbiome datasets.

## 2. Conceptual Models

We propose four conceptual models – the Union Model, the Intersection Model, the Gain Model, and the Loss Model – to describe the potential relationships between the HA microbiomes of hybrid individuals and those of their progenitors (see Fig. 1). In what follows, we delineate these four models, their underlying mechanisms, and their potential for impacting hybridization outcomes.

**Fig. 1.**
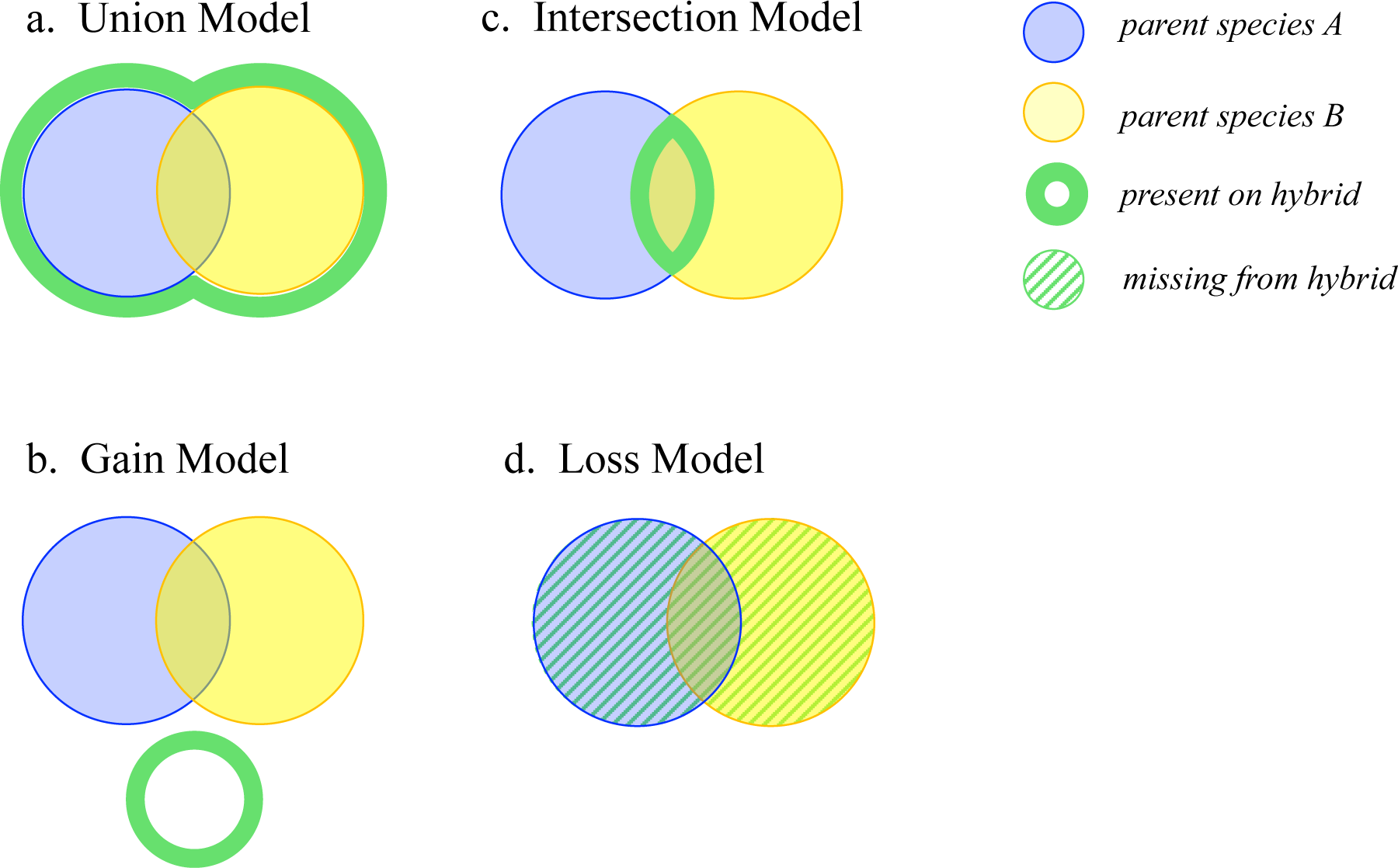
Four conceptual models for HA microbiomes of hybrid organisms. Hybrid organisms may host microbial taxa (a) found on *either* parent (union), (b) only found on *both* parents (intersection), or (c) found on *neither* parent (gain). Finally, (d) hybrids may be missing microbial taxa found on one or both parents (loss).

### Union Model

In this model, hybrid microbiomes include microbial taxa present on either progenitor. This could occur if carrying a particular host genome fosters colonization by associated microbial taxa. Notably, such fostering could emerge either directly through host interactions with the microbe (e.g., if specific hybrid and/or parent morphologies provide housing for symbiotic microbes^133–135^) or indirectly through effects on host behavior or ecology (e.g., if hybrids colonize a parental environment and subsequently acquire environmental microbes). To the extent that hybrid individuals share genetic material from both progenitors (note that this may vary depending on the extent of back-crossing), hybrids should support all microbes present on either parent. Said differently, in the Union Model, the host genome acts as a ‘ticket’ for acquiring a particular microbiome. Having two tickets (i.e., each representing a unique genomic component) results in the acquisition of two microbiomes, one from each progenitor.

Hybrids characterized by the Union Model should, in general, have more taxonomically diverse microbiomes than either progenitor. Importantly, greater taxonomic diversity could result in greater functional diversity as well^136^, with important consequences for host health and ecological performance. Consider a thought experiment wherein two different insect species are each limited to a distinct set of host plants based on the need for gut microbial detoxification of plant defensive compounds. If the hybrid offspring of these two insect species harbor the gut microbiomes of both parents, then hybrid microbiomes should be able to detoxify both sets of host plants, allowing hybrids to utilize all resources open to either progenitor. More broadly, greater functional capacity of hybrid microbiomes could enable hybrids to persist in habitats that are intermediate to their progenitors or across all habitats colonized by either progenitor. Beyond expanded function, a more diverse hybrid microbiome may have other benefits as well. Although a somewhat contentious theory, both in microbiome literature^137–139^ and more broadly, diversity-stability relationships^140, 141^ have long been posited to drive lower temporal variability and increased resistance to invasion in more diverse communities.^140^ If this is true for host-associated microbial communities, then hybrids following the Union Model may gain the advantage of having a more resilient microbiome that is more resistant to colonization by pathogens.^142^

However, there are likely costs to the Union Model as well. Most obvious are the challenges of bringing together large numbers of distinct microbial taxa from different progenitors. Consider Dobzhansky-Muller (DM) incompatibilities,^143–145^ which emerge in hybrid organisms due to mismatches between the genes from their two progenitors. If DM incompatibilities are a common outcome of combining different parental genomes, then analogous mismatches that result from combining different microbial metagenomes seem likely. Further, there could be mismatches between the microbial metagenome from one progenitor and the host genome from the other. Indeed, host-microbe and microbe-microbe incompatibilities are arguably even more likely than traditional DM incompatibilities simply because the microbial metagenome is typically much larger than the genome of the host.

### The Intersection Model

In this model hybrid microbiomes only include microbial taxa present on both progenitors. This could occur if a particular host genome hinders or prevents colonization by unassociated microbial taxa. Again, the underlying mechanism could be direct (e.g., changes in the host immune system) or indirect (e.g., changes in host behavior that alter exposure to environmental microbes). In either case, hybrids that carry genetic material from both progenitors will be more refractory to or isolated from a wider range of microbial taxa. Said differently, in the Intersection Model, each host genome acts as a ‘gate.’ Having two gates blocks a wider range of microbes, leaving only those taxa that are permitted access by both progenitors. Thus, hybrids characterized by the Intersection Model should have less taxonomically diverse microbiomes which could have consequences for functional diversity as well. For instance, in our previous insect example (see ‘Union Model’) the Intersection Model could leave hybrids without the ability to detoxify either set of parent host plants, placing a substantial limitation on feeding opportunities. This, in turn, could have impacts on fitness, leading to higher rates of starvation, underperformance due to toxin build-up, or even poisoning directly. Similar negative effects on survival could be possible due to more general mechanisms associated with microbial diversity as well, for example, the loss of microbiome stability and pathogen resistance. The benefit of the Intersection model, of course, is that it virtually eliminates opportunities for microbe-host or microbe-microbe incompatibilities. This is because, in the Intersection model, all microbe-host and microbe-microbe interactions that occur on the hybrid are already present on both progenitors.

### The Gain Model

In this model, hybrid microbiomes include microbial taxa not present on either progenitor. This is possible if HA microbiomes are idiosyncratically sensitive to specific gene combinations that arise from merging parent genomes. Broadly speaking, the Gain Model is the microbial equivalent of Bateson’s ‘saltational evolution’^146, 147^ or Goldschmidt’s ‘hopeful monsters’.^148, 149^ Like Bateson’s and Goldshmidt’s models, the Gain Model posits that hybridization can yield profound (saltational) changes in phenotype,^150, 151^ and that these phenotypic changes may enable hybrids to establish an entirely novel ecological niche relative to their progenitors.^148, 152, 153^ However, unlike Bateson and Goldschmidt, who focused on host genes, the Gain Model assumes that there are underlying microbial dimensions to the saltational change. Arguably, adding microbial dimensions provides even more opportunity for saltational change, again because of the vast size and diversity of functions encompassed by the microbial metagenome relative to the host genome itself. Once more, consider our hypothetical insect example (see ‘Union Model’). According to the Gain Model, hybrids should harbor an entirely new set of gut bacteria with novel taxa and potentially different detoxification properties as compared to their progenitors. Thus, rather than being able to use both of their parent’s host plants (Union Model), or neither of their parent’s host plants (Intersection Model), these ‘saltational’ hybrids could potentially colonize an entirely novel set of host plants not used by either progenitor.

More so than the Union, Intersection, or Loss Models, the Gain Model provides the building blocks for evolutionary innovation. This could result in rapid adaptation, escape from competition with their progenitors, or even reproductive isolation. Indeed, in extreme cases, the Gain Model may actually accelerate speciation.^152^ However, the Gain Model may have non- or maladaptive consequences as well. Notably, there is no *a priori* reason to believe that the acquisition of large numbers of novel microbial taxa will be generally beneficial to a host. In fact, there are many reasons to believe the opposite. In particular, the Gain Model describes a scenario of rapid evolutionary change (i.e., the introduction of novel microbial metagenomic content to the hybrid), that occurs far outside the confines of more typical host-microbe coevolutionary relationships forged over generations of symbiosis. As a result, the Gain Model exemplifies a “high risk, high reward” scenario, and novel microbes acquired by the hybrid could just as easily enhance or reduce host fitness. Thus, like Goldschmidt’s hopeful monsters, the Gain Model relies on ‘happy accidents,’^154^ meaning that many hybrid individuals are likely to fail for each ecological success.

### The Loss Model

In this model, hybrid microbiomes are missing microbial taxa that are present on one or both progenitors. While some loss – namely the loss of microbial taxa only present on one progenitor – is an inevitable part of the Intersection Model, The Loss Model includes the additional loss of microbial taxa present on both progenitors (note that defining the Loss Model to include loss from one or both parents is a necessary constraint of having a four dimensional index; splitting these two different forms of loss would require a higher dimensional index that could not be visualized). As in the Gain Model, the loss of microbes present on both progenitors describes a ‘saltational’ scenario that is possible if HA microbiomes are idiosyncratically sensitive to gene combinations of the progenitors. In contrast to the Gain Model, however, the saltational change invoked by the Loss Model is the deletion, rather than the addition of microbial taxa.

In general, the Loss Model should give rise to hybrid microbiomes with lower overall diversity and potentially lower functional capacity as well. Returning, for the last time, to our hypothetical insect complex, the Loss Model predicts that hybrid insects should lack microbes present on one and often both progenitors. In the case of the latter, hybrids would lose the ability to detoxify host plants that are usable by both of their progenitors. Like the Intersection Model, this could limit opportunities for feeding, cause toxin build-up or result in poisoning of hybrid insects. Lower microbiome diversity could also lead to a suite of additional challenges like greater microbiome instability and lower pathogen resistance. Again, however, the costs of low diversity microbiomes may be balanced out by the benefits of reducing opportunities for host-microbe or microbe-microbe incompatibilities. Indeed, in the limit of a hybrid organism fully characterized by the Loss Model, there would be no host-associated microbiome at all.

## 3. The 4H index and Quaternary Plots

To examine the importance of each of the four conceptual models to any given hybrid system, we introduce the 4H index, along with R package `HybridMicrobiomes` which can be used to calculate and graph the 4H index for any hybrid system. The 4H index uses the ‘core microbiomes’ of each host class (where we use ‘host class’ to refer to any one of the three types of hosts – the first progenitor, the second progenitor, or the hybrid – in a hybrid complex) to determine which microbial taxa are lost and gained on hybrid organisms relative to their progenitors. To define the core microbiome, we use a tunable parameter, ρ, which can range from ρ = 1 (microbial taxa are only considered if they are present on every host of a particular host class), to ρ = 0 (all microbial taxa are considered, regardless of the number of hosts they are found on). Consistent with the common definition of a core microbiome, we typically select higher values of ρ. This is based on the assumption that microbial taxa with strong consequences for host ecology and/or evolution should be detectable on the majority of hosts within a population. Notice, however, that researchers who have reason to suspect otherwise can use a lower value of ρ or can compare the 4H index across a range of ρ values (see Supplementary Information, Figs. S2.1 and S2.2 and Table S2). Thus, both the 4H index and the `HybridMicrobiomes` R package provide flexibility that can be decided within the context of a particular hybrid system (though a common set of parameters should be used for any comparison *between* hybrid systems). For the 4H index, ρ is the same across all host classes. However, each host class (i.e., each progenitor and the hybrid) is separately assigned its own core microbiome. Thus, a microbial taxon is part of a host’s core microbiome provided it is on at least !% hosts of that host class, where % is the number of hosts of each class and should be the same across all host classes (i.e., a balanced design with equal numbers of each progenitor and the hybrid; note that the `HybridMicrobiomes` package includes bootstrapping steps that will down sample datasets such that a balanced design is achieved).

For any given ρ, we define *P*_1_, *P*_2_ and *H* as the set of core microbial taxa present on the first progenitor, the second progenitor and hybrids respectively. Inspired by the Jaccard Index^155^ – a measure of species turnover between ecological communities – we then define the four dimensions of the 4H index (three independent dimensions) as the fraction of the total microbial diversity across the entire host species complex 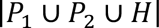) that is captured by each of our four conceptual models:

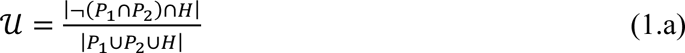

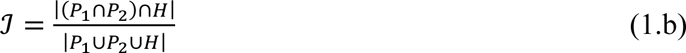

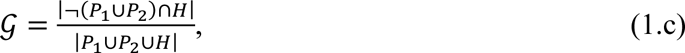

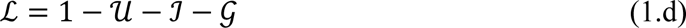

In equation (1), 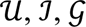 and 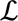 reflect the extent of Union, Intersection, Gain and Loss models respectively, and |*S*| denotes the cardinality of *S*, where *S* is any set. Briefly, 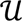 is the fraction of microbial taxa found on hybrids and on one but not both parents (note that this is a slight deviation from the conceptual Union Model, which does not distinguish between taxa found on one or both progenitors. This deviation is necessary to avoid double counting microbial taxa, and is further discussed below). 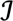 is the fraction of microbial taxa found on hybrids and on both progenitors. 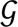 is the fraction of microbial taxa only found on hybrids and 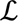 is the fraction of microbial taxa only found on progenitors. Function `FourHbootstrap` in `HybridMicrobiomes` takes as input any phyloseq object^156^, along with a vector specifying parent and hybrid classifications and calculates the 4H index over bootstrapped samples of hybrid organisms and their progenitors. Function `FourHcentroid` takes the output from `FourHbootstrap` and calculates the centroid of the bootstrapped samples. Finally, function `FourHcompare` takes the outputs from `FourHbootstrap` on multiple hybrid systems and uses a PERMANOVA test^157^ on the isometric log-ratio transformed^158^ 4H indices to determine whether different hybrid systems vary with respect to the importance of the Union, Intersection, Gain, and Loss Models respectively.

To visualize the 4H index (see Figure 2), which can be particularly helpful for comparison between systems, we introduce a quaternary plotting technique (i.e., a four-dimensional barycentric plot or an Aitchison Simplex^159^). This positions each of our four index dimensions (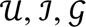 and 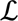) at a vertex of a triangular prism, with one edge of the prism connecting the Gain and Loss Models – henceforth termed the ‘transgressive axis’ – and the opposite edge connecting the Union and Intersection Models – henceforth termed the ‘parental axis.’ Function `FourHquaternary` takes the output from `FourHbootstrap` and generates an interactive quaternary plot of the bootstrapped samples with the option to include the centroid. Function `FourHquaternarycentroid` takes the output from `FourHbootstrap` and generates an interactive quaternary plot of only the centroids over the bootstrapped samples.

**Fig. 2.**
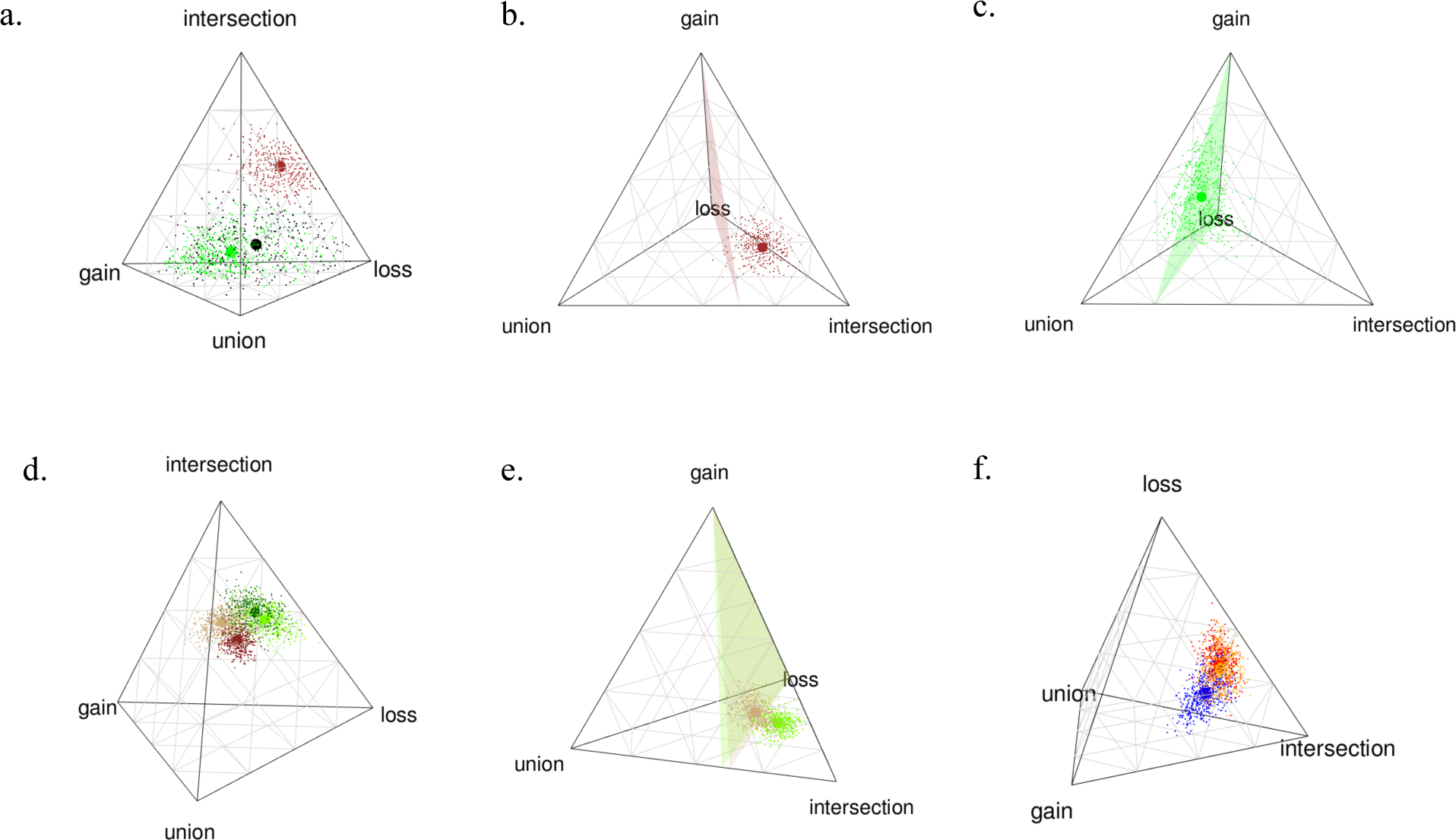
Quaternary plots showing 500 bootstrapped genus-level microbial samples (small circles) and the bootstrap centroid (large circles) of the 4H index for (a) gut microbiomes from hybrid *Kikihia* cicadas (black), *Neotoma* woodrats (brown) and *Aspidoscelis neomexicanus* whiptail lizards (green); (b,c) woodrat and lizard systems individually, along with the system null planes; (d) leaf (green) and rhizosphere (brown) bacterial/archaeal (16S rRNA, light) and fungal (ITS, dark) microbiomes from B73 line × Mo17 line maize hybrids; (e) B73 line × Mo17 line maize hybrid leaf and rhizosphere bacterial/archaeal systems, along with system null planes; (f) leaf bacterial/archaeal microbiomes from B73 line × Mo17 line (red), B73 line × CML103 line (yellow) and B73 line ×Mo18W line (blue) maize hybrids. For systems in (a-c), bootstraps consisted of 7 hybrid individuals and 7 of each progenitor. For systems in (d-f), bootstraps consisted of 10 hybrid individuals and 10 of each progenitor. A microbial genus was defined as being part of the core microbiome if at least 50% of hosts from a particular class carried that microbial genus.

As suggested above (see ‘Loss Model’), both our conceptual models and the four dimensions of the 4H index conflate microbial taxon loss due to the intersection of parental microbiomes (i.e., loss of microbial taxa only present on one progenitor) with broader microbial taxon loss (i.e., including loss of microbial taxa present on both progenitors). Thus, the 4H index does not indicate whether the microbial taxa that are lost versus retained by hybrid organisms represent microbial taxa that are shared by both progenitors or taxa that are only found on one progenitor. Unfortunately, conflation of these different types of loss is necessary for the 4H index to account for all microbial taxa in the system without double-counting microbial taxa (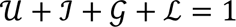) and while still using a maximum of four (beneficial for visualization) dimensions. To offset this constraint, and better identify the particular microbial taxa are lost by hybrid organisms, we develop ‘null planes.’ Specifically, we assume a null model wherein all microbial taxa present on progenitors are equally likely to be lost by hybrids. This allows us to define a plane bisecting the quaternary plot at the expected fraction of hybrid microbial taxa that should be shared with one versus both progenitors, assuming that there is no preferential loss of one over the other. For any value of 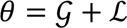 (i.e., the summed fractions of microbial taxa following the Gain and Loss Models), the null plane is given by:

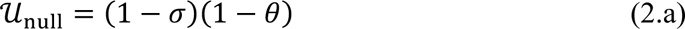

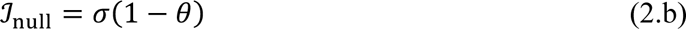

where 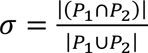 is the fraction of parental microbial taxa that are found on both progenitors. 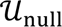 and 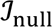 are thus the expected fractions of microbial taxa that should be found on only one parent versus both parents under null model assumptions. 4H-indices that lie more towards the 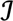 vertex relative to the null plane indicate that hybrids are disproportionately likely to retain microbes shared by both progenitors (i.e., loss is concentrated among microbial taxa only found on one of the two progenitors as in the Intersection Model). 4H-indices that lie more towards the 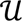 vertex relative to the null plane suggest that hybrids are disproportionately likely to retain microbes only found on one of the two progenitors (i.e., loss is concentrated among microbial taxa found on both progenitors and is, thus, saltational). By using the null plane as a reference, it is possible to assess the degree to which the loss occurs due to the intersection of parental microbiomes versus the broader ‘saltational’ loss of microbes present on both progenitors. Function `FourHnullplane` takes the output from `FourHbootstrap` and graphs the (average) null plane for a particular hybrid system onto a quaternary plot. Function `FourHplaneD` takes the output from `FourHbootstrap` and reports both the average distance between the expected, 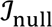_-_, and observed, 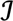, value of the intersection dimension, as well as the fraction, *p*, of bootstrap samples that lie further from the 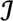 vertex than expected (this is useful for testing the hypothesis that microbes shared by both progenitors are more likely to be retained by the hybrid than microbes only found on one or other progenitors).

## 4. Example Analyses

Figure 2a shows quaternary plots (R version 4.2.1, phyloseq 1.41.1) comparing gut microbiomes from F_1_ crosses of *Neotoma* woodrats (brown)^122^, *Kikihia* cicadas with evidence of mitochondrial introgression (black)^160^, and a parthenogenetic *Aspidoscelis* lizard of hybrid origin (green, our own data). Table 1 shows the average values of the 4H indices for each system. Table 2 shows the average values of 5 (the fraction of the overall parental microbiome found on both progenitors), along with the mean distance between the predicted and observed value of 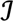, and the proportion of bootstrap samples that lie further from the 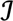 vertex than expected based on the null model. Despite the variation in life history (vertebrate vs. invertebrate, ectotherm vs. endotherm, herbivore vs. insectivore) and mode of hybridization (F_1_ crosses, mitochondrial introgression, hybrid speciation/parthenogenesis), the 4H index enables comparison across all systems. Hybrid woodrats, for example, are dominated by the Intersection Model (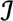 = 0.4927), while this is less important for hybrid cicadas (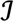 = 0.1337) and hybrid lizards (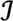 = 0.1160). Instead, lizards and cicadas feature a mix of the Gain and Loss models, with Gain being more important for lizards (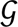 = 0.3438) and Loss being more important for cicadas (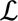 = 0.3758). Figure 2 (b,c) illustrate null planes for the woodrat and lizard systems. From the null planes and Table 2, we see that the two progenitor woodrat species share a larger fraction of their core microbes as compared to the two progenitor lizard species. Further, we see that hybrid woodrats are biased towards the intersection model as compared to the null plane (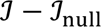 = 0.1358, *p* = 0.002). This means that hybrid woodrats are more likely to retain microbial taxa that are shared by both progenitors than they are to retain microbial taxa found on only one of the two progenitors. By contrast, hybrid lizards do not show any significant bias, being equally likely to retain (or lose) microbial taxa that are shared by both progenitors or only present on one of the two progenitors.

**Table 1.** Centroid values of the 4H index as calculated by the `FourHcentroid` function. Summing the values 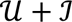 and 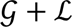 gives totals along the parental axis and the transgressive axis respectively, and can be used as a broader scale comparison between systems.

|  | Parental Axis |  |  | Transgressive Axis |  |  |
| --- | --- | --- | --- | --- | --- | --- |
| | $\mathcal{U}$ | $\mathcal{J}$ | $\mathcal{U} + \mathcal{J}$ | $\mathcal{G}$ | $\mathcal{L}$ | $\mathcal{G} + \mathcal{L}$ |
| *Neotoma* woodrat | 0.07296 | 0.4927 | 0.5657 | 0.0811 | 0.3532 | 0.4343 |
| *Aspidoscelis* lizard | 0.2941 | 0.1160 | 0.4101 | 0.3438 | 0.2461 | 0.5899 |
| *Kikihia* cicada | 0.1999 | 0.1337 | 0.3336 | 0.2906 | 0.3758 | 0.6664 |
| Maize B73×Mo17<br>(leaf, bacteria) | 0.1272 | 0.4949 | 0.6221 | 0.0575 | 0.3205 | 0.3780 |
| Maize B73×Mo17<br>(rhizosphere, bacteria) | 0.1892 | 0.5000 | 0.6892 | 0.1624 | 0.1484 | 0.3108 |
| Maize B73×Mo17<br>(leaf, fungi) | 0.1029 | 0.5238 | 0.6267 | 0.0902 | 0.2832 | 0.3734 |
| Maize B73×Mo17<br>(rhizosphere, fungi) | 0.1936 | 0.4226 | 0.6162 | 0.1524 | 0.2314 | 0.3838 |
| Maize B73×CML103<br>(leaf, bacteria) | 0.1070 | 0.5044 | 0.6114 | 0.0798 | 0.3087 | 0.3885 |
| Maize B73×Mo18W<br>(leaf, bacteria) | 0.1601 | 0.4784 | 0.6385 | 0.1632 | 0.1983 | 0.3615 |
| Maize B73×CML103<br>(rhizosphere, fungi) | 0.2152 | 0.4468 | 0.6620 | 0.1123 | 0.2256 | 0.3379 |
| Maize B73×Mo18W<br>(rhizosphere, fungi) | 0.1433 | 0.4635 | 0.6068 | 0.1146 | 0.2785 | 0.3931 |

**Table 2.** Fraction of shared microbial taxa among progenitors, 5, as calculated by the `FourHcentroid` function, as well as average displacement from the null plane and proportion of bootstrap samples falling further from the 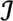 vertex than the null plane, both as calculated by the `FourHnullplaneD` function.

| | $\sigma$ | $\mathcal{I} - \mathcal{I}_{\text{null}}$ | $p$ |
| --- | --- | --- | --- |
| *Neotoma* woodrat | 0.6273 | 0.1358 | 0.002 |
| *Aspidoscelis* lizard | 0.2490 | 0.0110 | 0.364 |
| *Kikihia* cicada | 0.2827 | 0.0373 | 0.178 |
| Maize B73×Mo17<br>(leaf, bacteria) | 0.5984 | 0.1208 | 0 |
| Maize B73×Mo17<br>(rhizosphere, bacteria) | 0.6291 | 0.0663 | 0 |
| Maize B73×Mo17<br>(leaf, fungi) | 0.6483 | 0.1167 | 0 |
| Maize B73×Mo17<br>(rhizosphere, fungi) | 0.5664 | 0.0742 | 0 |
| Maize B73×CML103<br>(leaf, bacteria) | 0.6475 | 0.1077 | 0 |
| Maize B73×Mo18W<br>(leaf,bacteria) | 0.6121 | 0.0872 | 0 |
| Maize B73×CML103<br>(rhizosphere, fungi) | 0.5565 | 0.0781 | 0 |
| Maize B73×Mo18W<br>(rhizosphere, fungi) | 0.6145 | 0.0910 | 0 |

Figure 2d shows quaternary plots of both the phyllosphere (green) and rhizosphere (brown) of hybrid maize (B73 line × Mo17 line) for both bacterial/archaeal (16S rRNA gene, light shade) and fungal (ITS1 gene, dark shade) microbiomes.^125^ Figure 2e shows the same bacterial/archaeal microbiomes but includes their respective null planes, and Figure 2f compares the bacterial/archaeal phyllosphere microbiomes across three different maize hybrids: B73 line × Mo17 line (red; stiff stalk vs. non-stiff stalk varieties^125^), B73 line × CML103 line (yellow; temperate vs. tropical varieties^161^), and B73 line × Mo18W line (blue; flooding sensitive vs. flooding insensitive^162^). Tables 1 and 2 show the corresponding values of the 4H indices, as well as relationships of the hybrid microbiomes to their respective null planes. As with our animal examples, our analysis of maize hybrids demonstrates the versatility of the 4H index, and how the 4H index can be used to compare not only between microbiomes from different host species, but also between microbiomes from different parts of a single organism (roots vs. leaf), or different microbial taxonomic groups (bacteria vs. fungi). From the maize comparisons, we see that the entire maize system is dominated by the Intersection Model. However, in general, the hybrid rhizosphere (brown) is more prone to Union and Gain of microbial taxa, whereas the hybrid phyllosphere is more prone to Loss.

One of the benefits of the 4H index is the fact that it can be applied to any hybrid system, regardless of the type of host, the type of microbiome, microbiome composition, or even microbiome diversity. This flexibility follows from the fact that the 4H index is monotonic with respect to each vertex/dimension 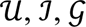 and 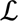), density invariant, and replication invariant (see Supplemental Information).^163^ Despite this, some standardization of datasets from different systems is necessary for fair comparison. For example, although the 4H index can be applied to microbiomes at any taxonomic scale, higher taxonomic scales predict a greater importance of the Intersection Model (e.g., hybrids are more likely to share microbial phyla with progenitors than they are to share microbial ASVs; see Supplemental Information Fig. S1.1, S1.2, and Table S.1). Thus, systems should always be compared using the same taxonomic scale. Likewise, defining the core microbiome based on a lower fraction of hosts also favors intersection (at least some hybrids and some of each parental species are likely to have a particular microbial taxon, even if it is just a transient acquisition from the environment; see Supplemental Information Fig. S.2.1, S.2.2, and Table S.2). Again, then, it is important to use the same value of ρ for all systems that are being compared. Host sample size has a smaller, but still detectable effect, resulting in somewhat different trends across systems, but generally appearing to shift the 4H index towards the parental axis and away from the transgressive axis (see Supplemental Information Fig. S3.1, S.3.2, and Table S.3). Although the effect of host sample size is relatively small, particularly for larger host sample sizes, it is still best to compare systems by subsampling to the smallest number of hosts available for any host class across all systems (e.g., see Fig. 2 where we were limited to 7 individuals based on the number of available cicada microbiomes). Finally, sequencing depth has almost no impact on predictions, at least for >1000 reads or more. This last feature of the 4H index is a benefit of focusing on core microbiomes since low abundance microbial taxa that are likely to be missed at low read depths are unlikely to be part of the core of any given species. For this reason, it is largely unnecessary to standardize for read depth across systems. Nevertheless, function `FourHbootstrap` does have the option to rarefy microbiome samples to a lower read depth than the minimal number of reads of the lowest sample. This allows for standardization of read depth across systems, if desired.

## 5. Conclusions

The advent of low-cost sequencing has greatly contributed to our understanding of the prevalence and importance of both hybridization and host-associated microbiomes on host ecological traits and evolutionary consequences. These two fields come together in the study of host-associated microbiomes of hybrid organisms – a newly emerging area of research across disciplines ranging from agricultural science to ecology and conservation. In this paper, we integrate four conceptual models to develop a framework for understanding the relationship between hybrid microbiomes and the microbiomes of their progenitors. We then use these models to develop a four-dimensional (three independent dimensions) metric – the 4H index – to describe where a particular hybrid complex falls among our four models. Importantly, the 4H index facilitates comparisons across widely disparate systems, ultimately making it possible to identify patterns that emerge across hybrid microbiomes from different organisms. For example, the 4H index could be used to determine whether there are systematic differences between hybrid plant versus hybrid animal microbiomes, or between hybrid vertebrate versus hybrid invertebrate microbiomes. Likewise, the 4H index could be used to determine how phylogenetic and/or phenotypic distances between progenitors or ploidy level impact the hybrid microbiome.

While we envision the 4H index primarily as a tool for analyzing hybrid microbiomes, it is worth noting that this same framework can be applied to any triplet of host species, where one of the three host species is in some way ‘intermediate’ to the other two. Thus, for example, a 4H index could be calculated for the microbiomes of organisms from ecotonal habitat, and then compared to the microbiomes of organisms from the two pure habitat types on either end of the ecotone^164^, even if it is the same host taxon across the entire zone. Likewise, a 4H index could be calculated for species (e.g., swordtail males, *Xiphophorus nigrensis*) that exhibit three discrete size classes, with one size class being intermediate to the other two.^165^ Similarly, a 4H index could be calculated for captive animals fed two different pure diets as compared to captive animals fed a mixed diet. In these scenarios, the interpretation of our four conceptual models would change. However, because the 4H-metric is defined solely based on distributions of microbial presence/absence across non-overlapping sets of host classes, it is valid for any analysis where there is ecological, evolutionary, morphological, or physiological reason to believe that one host class falls between the other two host classes.

