## SupplementaryInformation for "A Conceptual Framework for Host-Associated Microbiomes of Hybrid Organisms"

**Supplementary Information**

**S1. Example Datasets**

*Neotoma* woodrats and *Kikihia* cicadas: Details regarding hybrid system, sample collection and sample processing can be found in the respective publications.[122,160] For both systems, we downloaded FASTQ files from the NCBI sequence read archive under Bioprojects PRJNA887535 and PRJNA879614 respectively. We then used the QIIME 2 v.2022–8 software[166] to process contigs and perform taxonomic attribution. Briefly, contig sequences were imported into QIIME2 via the *qiime tools import* command using a manifest file and input format PairedEndFastqManifestPhred33. Quality control of contigs was performed using the *qiime quality-filter* command, and reads were trimmed to 250 bp using the *qiime deblur denoise-16S* command. For taxonomic attribution, we used the silva database with the *silva-138-99-515-806-nb-classifier.qza*. Finally, taxonomic classifications were pooled to the genus-level using the *qiime taxa collapse* command at level 6. All analyses for these systems were performed on the resulting OTU tables at the genus-level.

*Aspidoscelis* lizards: *Aspidoscelis inornatus*, *A. marmoratus* and *A. neomexicanus* were captured in two locations in Soccorro County, NM, USA. *A. inornatus* and *A. neomexicanus* were syntopic at one location, and *A. marmoratus* and *A. neomexicanus* were syntopic at the other. Upon capture, 'gut' microbiomes were collected from individual lizards by inserting a sterile swab (Puritan HydraFlock #253318H; tip dimensions 0.7 cm length × 0.3556 cm [0.14 in.] diameter) into the cloaca while gently rotating and pulsing it along the anterior-posterior axis of the animal. Swabs were immediately deposited in DNA/RNA Shield (Zymo R1109) and were then sent to Zymo Research for processing by their ZymoBIOMICS service. Zymo provides a range of OTU tables across varying taxonomic levels from amplicon sequence variants (ASVs) to phyla. Before analyzing the ASV table, we removed all sequences that could not be mapped to the bacterial domain. Before analyzing the tables for other taxonomic levels, we removed the row that was not mapped to either Bacteria or Archaea. Following these modifications, ttables were used without further processing for subsequent analysis.

Maize: Details regarding hybrid system, sample collection and sample processing can be found in the corresponding publication.[125] We downloaded fungal and bacterial/archaeal ASV tables directly from the supplementary material provided by Wagner et al. (2020)[125] and used these ASV tables for all future analyses. Briefly, we converted fungal and bacterial/archaeal ASV tables to biom files using the `write_biom` function from the `biomformat` package in R (R 4.2.1).[167] We then imported the biom files into QIIME2 using the *qiime tools import* command. Next, we pooled microbial taxa to various taxonomic levels from genus through to phylum using the *qiime taxa collapse* command from levels 2 through 6. We then used the resulting OTU tables (along with the ASV table) for all subsequent analyses.

**S2. Monotonicity, Density Invariance and Replication Invariance**

Monotonicity: Our definition of monotonicity follows from an analogous definition used to validate measures of β-diversity.[163] That is, separately adding microbial taxa that follow different conceptual models to each of two systems with initially identical 4H indices should result in divergence of their 4H indices. Suppose, for example, that we have two systems (i.e., two hybrid complexes) with identical 4H indices. If we add $u$ microbial taxa that follow the Union Model to the first system, and $i$ microbial taxa that follow the Intersection Model to the second system, then this should result in the 4H indices of the two systems becoming further apart. Define the original indices for the first and second systems as:

$$\mathcal{U}_0 = \frac{U}{T}, \qquad \mathcal{I}_0 = \frac{I}{T}, \qquad \mathcal{G}_0 = \frac{G}{T}, \qquad \mathcal{L}_0 = 1 - \mathcal{U}_0 - \mathcal{I}_0 - \mathcal{G}_0 \tag{S1}$$

Further define the new index for the first system as:

$$\mathcal{U}_1 = \frac{U+u}{T+u}, \quad \mathcal{I}_1 = \frac{I}{T+u}, \qquad \mathcal{G}_1 = \frac{G}{T+u}, \qquad \mathcal{L}_1 = 1 - \mathcal{U}_1 - \mathcal{I}_1 - \mathcal{G}_1 \tag{S2}$$

and the new index for the second system as:

$$\mathcal{U}_2 = \frac{U}{T+i}, \qquad \mathcal{I}_2 = \frac{I+i}{T+i}, \qquad \mathcal{G}_2 = \frac{G}{T+i}, \qquad \mathcal{L}_2 = 1 - \mathcal{U}_2 - \mathcal{I}_2 - \mathcal{G}_2 \tag{S3}$$

In equations (S1-S3), $U$ is the number of microbial taxa that follow the Union Model in each of the original systems, $I$ is the number of microbial taxa that follow the Intersection Model in each of the original systems, $G$ is the number of microbial taxa that follow the Gain Model in each of the original systems, and $T$ is the total number of microbial taxa in each of the original systems (see equation 1 in the Main Text). The Euclidean distance between the original first and second systems is zero (they are identical). However, after addition of the $u$ taxa to the first system and the $i$ taxa to the second system, the Euclidean distance becomes:

$$d = \sqrt{\left(\frac{U+u}{T+u} - \frac{U}{T+x}\right)^2 + \left(\frac{I}{T+u} - \frac{I+x}{T+x}\right)^2 + \left(\frac{G}{T+u} - \frac{G}{T+x}\right)^2 + \left(\frac{L}{T+u} - \frac{L}{T+x}\right)^2} \tag{S4}$$

Notice that, because the expression under the square root is a sum of squares, equation (S4) will always be greater than or equal to zero, and will be strictly greater than zero provided that at least one term in the sum is positive. The requirement that every term in the sum be zero is:

$$x = -\frac{u(T-U)}{U+u} \qquad \text{and} \qquad x = -\frac{Iu}{T-I+u} \qquad \text{and} \qquad x \neq u \tag{S5}$$

Since equation (S5) will only be true for $x = 0, -T$, the distance between the two systems increases for all $x, u > 0$ and the monotonicity condition is fulfilled. Notice that this demonstration of monotonicity is identical regardless of the vertices ($\mathcal{U}$, $\mathcal{I}$, $\mathcal{G}$, and $\mathcal{L}$) selected for differentiating systems. Thus, we have only shown one example using $\mathcal{U}$ and $\mathcal{I}$.

Density Invariance: Again, our definition of density invariance follows from analogous definitions for β-diversity.[163] For β-diversity, a metric that is density invariant should not be sensitive to the raw abundances of species in each assemblage. The 4H index is invariant to the absolute number of reads of any given microbial taxon on any given host, both because the 4H index is based on presence/absence data and because it uses rarefied microbiome datasets. Additionally, however, the 4H index is invariant to the absolute number of hosts that harbor any particular microbial taxon. This is a result of our definition of the core microbiome, which is based on the fraction of hosts carrying a particular taxon, rather than the absolute number of hosts carrying that taxon. Further, and perhaps most importantly, particularly for comparisons across hybrid systems, the 4H index is invariant to the absolute diversity of the microbiomes of any given hybrid system. Thus, adding a proportional number of microbial taxa to the $\mathcal{U}$, $\mathcal{I}$, $\mathcal{G}$, and $\mathcal{L}$ categories of each host class does not change the overall value of the 4H index. To demonstrate this, we define the original index of a system as:

$$\mathcal{U}_0 = \frac{U}{T}, \qquad \mathcal{I}_0 = \frac{I}{T}, \qquad \mathcal{G}_0 = \frac{G}{T}, \qquad \mathcal{L}_0 = \frac{L}{T} \tag{S6}$$

Next, define the new index of the system as:

$$\mathcal{U}_1 = \frac{U+\omega\frac{U}{T}}{T+\omega\left(\frac{U}{T}+\frac{I}{T}+\frac{G}{T}+\frac{L}{T}\right)}, \mathcal{I}_1 = \frac{I+\omega\frac{I}{T}}{T+\omega\left(\frac{U}{T}+\frac{I}{T}+\frac{G}{T}+\frac{L}{T}\right)}, \mathcal{G}_1 = \frac{G+\omega\frac{G}{T}}{T+\omega\left(\frac{U}{T}+\frac{I}{T}+\frac{G}{T}+\frac{L}{T}\right)}, \mathcal{L}_1 = \frac{L+\omega\frac{L}{T}}{T+\omega\left(\frac{U}{T}+\frac{I}{T}+\frac{G}{T}+\frac{L}{T}\right)} \tag{S7}$$

where we have added taxa to each model proportional to their existing percentages across the hybrid system. The distance between the first index in equation (S6) and second index in equation (S7) is:

$$d = \sqrt{\left(\frac{U}{T} - \frac{U + \omega\frac{U}{T}}{T + \omega\left(\frac{U}{T} + \frac{I}{T} + \frac{G}{T} + \mathcal{L}_0\right)}\right)^2 + \left(\frac{I}{T} - \frac{I + \omega\frac{I}{T}}{T + \omega\left(\frac{U}{T} + \frac{I}{T} + \frac{G}{T} + \frac{L}{T}\right)}\right)^2 + \left(\frac{G}{T} - \frac{G + \omega\frac{G}{T}}{T + \omega\left(\frac{U}{T} + \frac{I}{T} + \frac{G}{T} + \frac{L}{T}\right)}\right)^2 + \left(\frac{L}{T} - \frac{L + \omega\frac{L}{T}}{T + \omega\left(\frac{U}{T} + \frac{I}{T} + \frac{G}{T} + \frac{L}{T}\right)}\right)^2} \quad \text{(S8)}$$

which simplifies to $d = 0$. Thus, the system is density invariant with respect to the absolute diversity of HA microbiomes across the hybrid complex.

Replication Invariance: As with the previous two criteria, our definition of replication invariance follows from an analogous definition for β-diversity.[163] Specifically, the 4H index should be invariant to pooling of samples with identical 4H indices. To demonstrate that this is the case, consider two systems. We define the indices for the first system as:

$$\mathcal{U}_1 = \frac{U_1}{T_1}, \qquad \mathcal{I}_1 = \frac{I_1}{T_1}, \qquad \mathcal{G}_1 = \frac{G_1}{T_1}, \qquad \mathcal{L}_1 = \frac{L_1}{T_1} \quad \text{(S9)}$$

and the indices for the second system as:

$$\mathcal{U}_2 = \frac{U_2}{T_2}, \qquad \mathcal{I}_2 = \frac{I_2}{T_2}, \qquad \mathcal{G}_2 = \frac{G_2}{T_2}, \qquad \mathcal{L}_2 = \frac{L_2}{T_2} \quad \text{(S10)}$$

with the constraint that $\frac{U_2}{T_2} = \frac{U_1}{T_1}, \frac{I_1}{T_1} = \frac{I_2}{T_2}, \frac{G_1}{T_1} = \frac{G_2}{T_2}$ and $\frac{L_1}{T_1} = \frac{L_2}{T_2}$ (i.e., the two systems have identical 4H indices prior to pooling). In this case, the new 4H index for the pooled system will be

$$\mathcal{U}_p = \frac{U_1+U_2}{T_1+T_2} = \frac{2U_1}{2T_1}, \quad \mathcal{I}_p = \frac{I_1+I_2}{T_1+T_2} = \frac{2I_1}{2T_1}, \qquad \mathcal{G}_p = \frac{G_1+G_2}{T_1+T_2} = \frac{2G_1}{2T_1}, \qquad \mathcal{L}_p = \frac{L_1+L_2}{T_1+T_2} = \frac{2L_1}{2T_1} \quad \text{(S11)}$$

which clearly reduces to equation (S9), demonstrating replication invariance.

### S3. Additional Figures and Tables

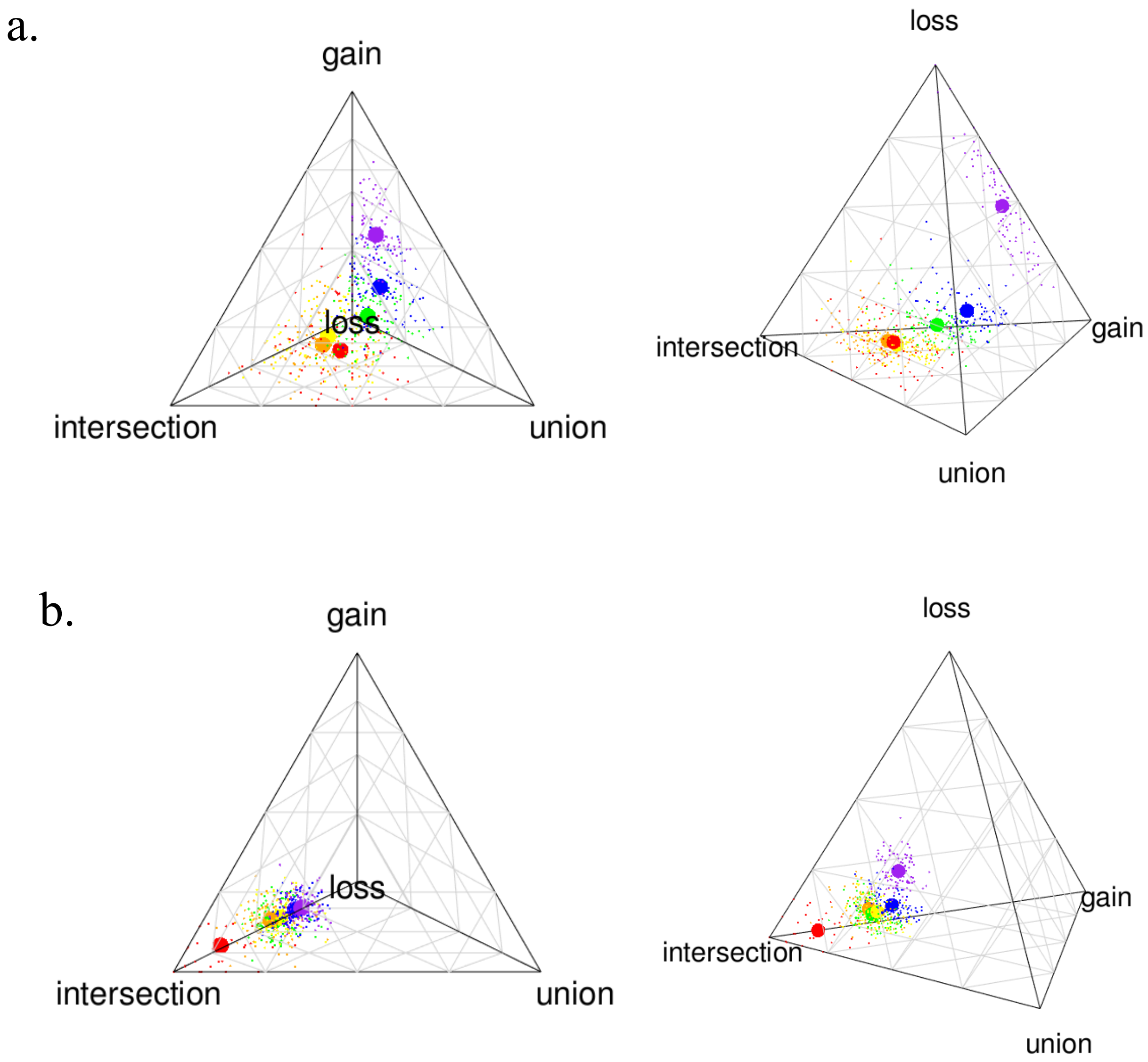

**Fig. S1.1** Quaternary plot showing 100 bootstrapped samples (small circles) and the bootstrap centroids (large circles) of the 4H index for gut microbiomes from (a) *Aspidoscelis neomexicanus* lizards and (b) B73 line × Mo17 line maize crosses. In quaternary plots, we show the 4H index for microbial taxa pooled by phylum (red), class (orange), order (yellow), family (green), genus (blue) and amplicon sequence variant (ASV, purple). Values of the corresponding centroids and total sums for the parental and transgressive axes are shown, below, in Table S.1. For both systems, we used $\rho = 0.5$ and subsampled 12 individuals of each progenitor and 12 hybrids.

**Table S.1** Variation in the 4H index as a function of the microbial taxonomic scale considered. Values are based on 100 bootstrapped samples of 12 individuals of each progenitor and 12 individuals of the hybrid, assuming $\rho = 0.5$ and using the FourHbootstrap and FourHcentroid functions from the HybridMicrobiomes package in R.

| | Parental Axis | | | Transgressive Axis | | |
|---|---|---|---|---|---|---|
| *Aspidoscelis* lizards (16S rRNA gene, cloacal swabs) | | | | | | |
| Taxonomic Scale | $\mathcal{U}$ | $\mathcal{I}$ | $\mathcal{U}+\mathcal{I}$ | $\mathcal{G}$ | $\mathcal{L}$ | $\mathcal{G}+\mathcal{L}$ |
| Phylum | 0.3504 | 0.4145 | 0.7649 | 0.1563 | 0.0788 | 0.2351 |
| Class | 0.2990 | 0.4595 | 0.7585 | 0.1761 | 0.0654 | 0.2415 |
| Order | 0.3054 | 0.4383 | 0.7437 | 0.2047 | 0.0515 | 0.2562 |
| Family | 0.3495 | 0.2666 | 0.6161 | 0.2461 | 0.1378 | 0.3839 |
| Genus | 0.3225 | 0.1758 | 0.4983 | 0.3240 | 0.1778 | 0.5018 |
| ASV | 0.1169 | 0 | 0.1169 | 0.3714 | 0.5116 | 0.883 |
| B73 line × Mo17 line maize (16S rRNA gene; rhizosphere) | | | | | | |
| Taxonomic Scale | $\mathcal{U}$ | $\mathcal{I}$ | $\mathcal{U}+\mathcal{I}$ | $\mathcal{G}$ | $\mathcal{L}$ | $\mathcal{G}+\mathcal{L}$ |
| Phylum | 0.0778 | 0.8174 | 0.8952 | 0.0755 | 0.0293 | 0.1048 |
| Class | 0.1461 | 0.6083 | 0.7544 | 0.1332 | 0.1124 | 0.2456 |
| Order | 0.1703 | 0.5811 | 0.7514 | 0.1434 | 0.1052 | 0.2486 |
| Family | 0.1639 | 0.6023 | 0.7662 | 0.1382 | 0.0955 | 0.2337 |
| Genus | 0.1894 | 0.5209 | 0.7103 | 0.1599 | 0.1298 | 0.2897 |
| ASV | 0.1629 | 0.4518 | 0.6147 | 0.1336 | 0.2517 | 0.3853 |

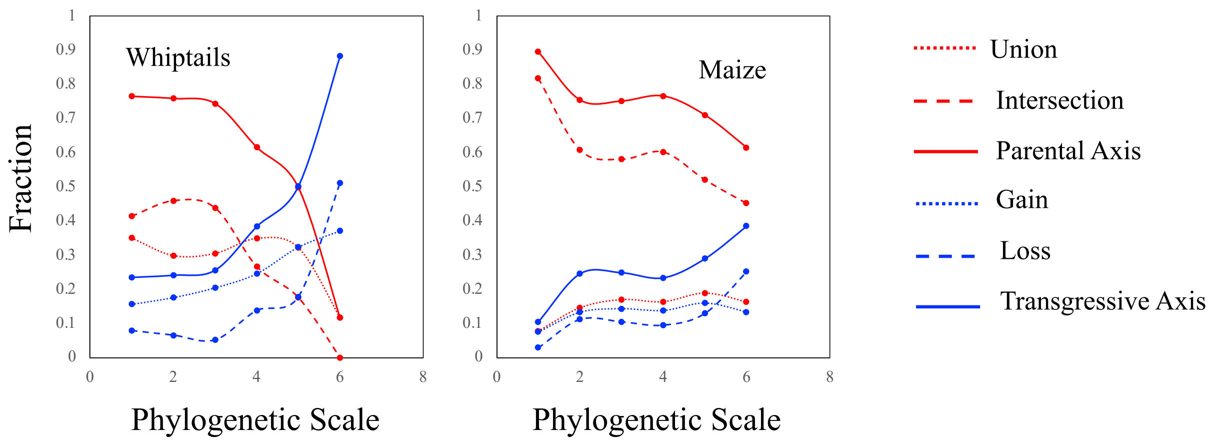

**Fig. S.1.2.** Variation in the 4H index as a function of the microbial taxonomic scale considered (1 = phylum, 2 = class, 3 = order, 4 = family, 5 = genus, 6 = ASV, See Table S.1).

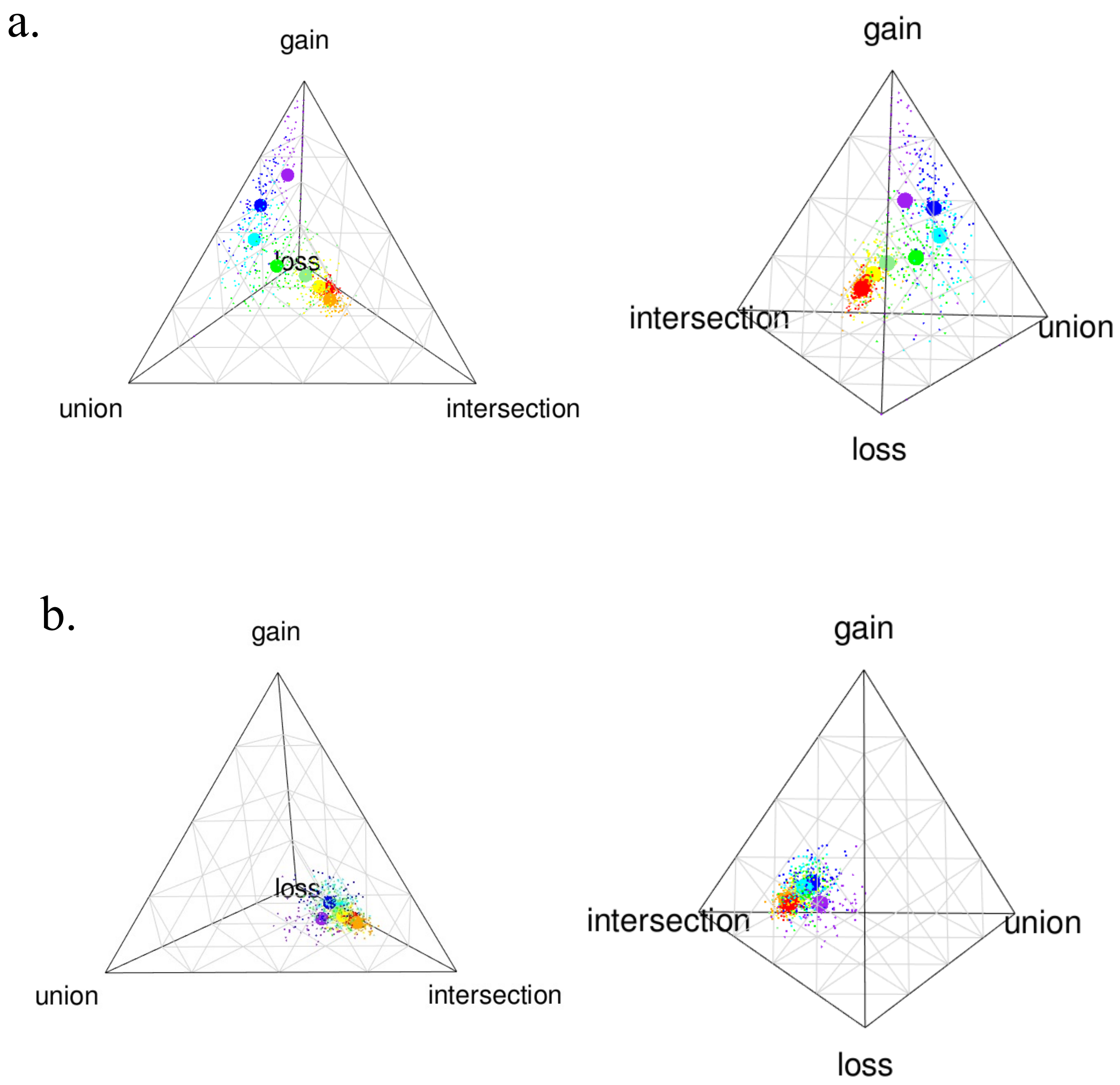

**Fig. S2.1** Quaternary plot showing 100 genus-level bootstrapped samples (small circles) and the bootstrap centroids (large circles) of the 4H index for gut microbiomes from (a) *Aspidoscelis neomexicanus* lizards and (b) B73 line × Mo17 line maize crosses. In all panels, we show $\rho$ = 0.1 (red), $\rho$ = 0.2, (orange), $\rho$ = 0.3 (yellow), $\rho$ = 0.4 (light green), $\rho$ = 0.5 (green), $\rho$ = 0.6 (cyan), $\rho$ = 0.7 (blue), and $\rho$ = 0.8 (purple). Values of the corresponding centroids and total sums for the parental and transgressive axes are shown, below, in Table S.2. For both systems, subsampled 12 individuals of each progenitor and 12 hybrids.

**Table S.2** Variation in the 4H index as a function of the definition of the core microbiome. Values are based on 100 bootstrapped samples of 12 individuals of each progenitor and 12 individuals of the hybrid, pooling microbial taxa to genus and using the `FourHbootstrap` and `FourHcentroid` functions from the `HybridMicrobiomes` package in R.

| | Parental Axis | | | Transgressive Axis | | |
|---|---|---|---|---|---|---|
| *Aspidoscelis* lizards (16S rRNA gene, cloacal swabs) | | | | | | |
| $\rho$ | $\mathcal{U}$ | $\mathcal{I}$ | $\mathcal{U}+\mathcal{I}$ | $\mathcal{G}$ | $\mathcal{L}$ | $\mathcal{G}+\mathcal{L}$ |
| 0.1 | 0.1833 | 0.3500 | 0.5333 | 0.2039 | 0.2628 | 0.4667 |
| 0.2 | 0.2209 | 0.3881 | 0.609 | 0.1916 | 0.1993 | 0.3909 |
| 0.3 | 0.2334 | 0.3357 | 0.5691 | 0.2420 | 0.1889 | 0.4309 |
| 0.4 | 0.2595 | 0.2788 | 0.5383 | 0.2849 | 0.1768 | 0.4617 |
| 0.5 | 0.3264 | 0.1735 | 0.4999 | 0.3140 | 0.1861 | 0.5001 |
| 0.6 | 0.3490 | 0.0601 | 0.4091 | 0.4061 | 0.1847 | 0.5908 |
| 0.7 | 0.2663 | 0.0145 | 0.2808 | 0.5187 | 0.2005 | 0.7192 |
| 0.8 | 0.0890 | 0 | 0.089 | 0.5820 | 0.3290 | 0.911 |
| B73 line × Mo17 line maize (16S rRNA gene; rhizosphere) | | | | | | |
| $\rho$ | $\mathcal{U}$ | $\mathcal{I}$ | $\mathcal{U}+\mathcal{I}$ | $\mathcal{G}$ | $\mathcal{L}$ | $\mathcal{G}+\mathcal{L}$ |
| 0.1 | 0.1470 | 0.5610 | 0.708 | 0.1195 | 0.1724 | 0.2919 |
| 0.2 | 0.1530 | 0.5925 | 0.7455 | 0.1167 | 0.1377 | 0.2544 |
| 0.3 | 0.1825 | 0.5411 | 0.7236 | 0.1392 | 0.1372 | 0.2764 |
| 0.4 | 0.1884 | 0.5208 | 0.7092 | 0.1549 | 0.1359 | 0.2908 |
| 0.5 | 0.1870 | 0.5167 | 0.7037 | 0.1594 | 0.1369 | 0.2963 |
| 0.6 | 0.1806 | 0.5106 | 0.6912 | 0.1741 | 0.1347 | 0.3088 |
| 0.7 | 0.2023 | 0.4854 | 0.6877 | 0.1882 | 0.1241 | 0.3123 |
| 0.8 | 0.2350 | 0.4683 | 0.7033 | 0.1179 | 0.1788 | 0.2967 |

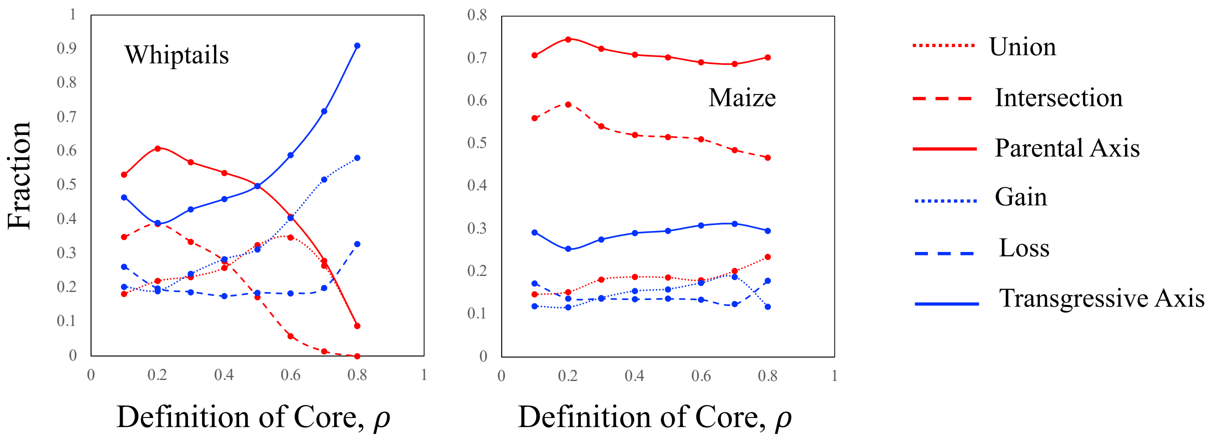

**Fig. S.2.2.** Variation in the 4H index as a function of the definition of the core microbiome (See Table S.2).

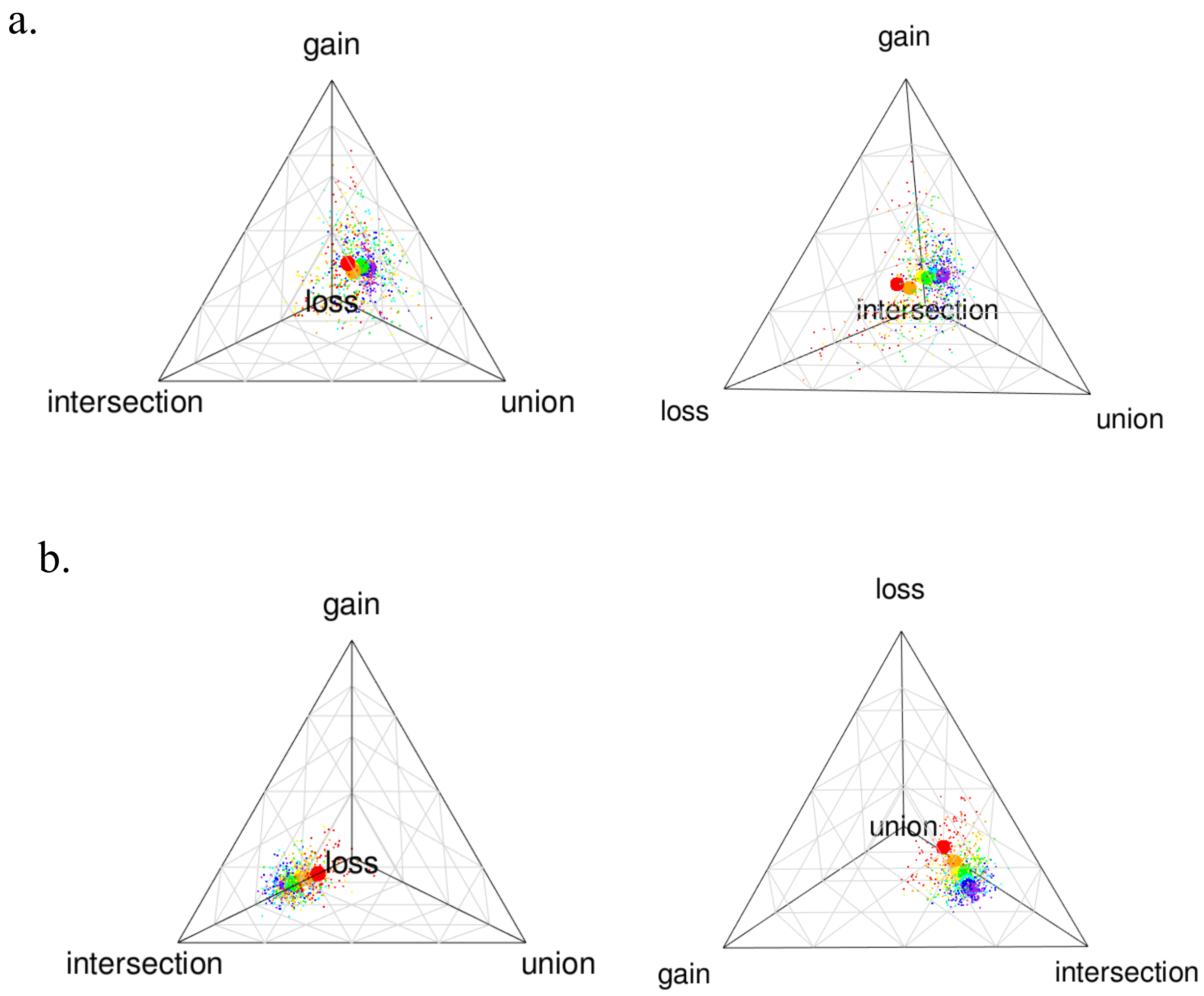

**Fig. S3.1** Quaternary plot showing 100 genus-level bootstrapped samples (small circles) and the bootstrap centroids (large circles) of the 4H index for (a) gut microbiomes from *Aspidoscelis neomexicanus* lizards and (b) B73 line × Mo17 line maize crosses (b). In all panels, we show bootstraps of 4 (red), 6 (orange), 8 (yellow), 10 (green), 12 (cyan), 14 (blue) and 16 (purple) individuals of each host class. Values of the corresponding centroids and total sums for the parental and transgressive axes are shown, below, in Table S.3. For both systems, we used $\rho = 0.5$.

**Table S.3** Variation in the 4H index as a function of the number of hosts per bootstrap sample. Values are based on 100 genus-level bootstrapped samples assuming $\rho = 0.5$ and using the FourHbootstrap and FourHcentroid functions from the HybridMicrobiomes package in R.

| | Parental Axis | | | Transgressive Axis | | |
|---|---|---|---|---|---|---|
| *Aspidoscelis* lizards (16S rRNA gene, cloacal swabs) | | | | | | |
| $N$ | $\mathcal{U}$ | $\mathcal{I}$ | $\mathcal{U} + \mathcal{I}$ | $\mathcal{G}$ | $\mathcal{L}$ | $\mathcal{G} + \mathcal{L}$ |
| 4 | 0.2415 | 0.1559 | 0.3974 | 0.2981 | 0.3044 | 0.6025 |
| 6 | 0.2798 | 0.1638 | 0.4436 | 0.2839 | 0.2726 | 0.5565 |
| 8 | 0.2901 | 0.1824 | 0.4725 | 0.3163 | 0.2111 | 0.5274 |
| 10 | 0.3174 | 0.1581 | 0.4755 | 0.3182 | 0.2062 | 0.5244 |
| 12 | 0.3277 | 0.1612 | 0.4889 | 0.3307 | 0.1805 | 0.5112 |
| 14 | 0.3400 | 0.1639 | 0.5039 | 0.3204 | 0.1756 | 0.496 |
| 16 | 0.3584 | 0.1527 | 0.5111 | 0.3275 | 0.1614 | 0.4889 |
| B73 line × Mo17 line maize (16S rRNA gene; rhizosphere) | | | | | | |
| $N$ | $\mathcal{U}$ | $\mathcal{I}$ | $\mathcal{U} + \mathcal{I}$ | $\mathcal{G}$ | $\mathcal{L}$ | $\mathcal{G} + \mathcal{L}$ |
| 4 | 0.2087 | 0.3928 | 0.6015 | 0.1654 | 0.2332 | 0.3986 |
| 6 | 0.1935 | 0.4535 | 0.647 | 0.1621 | 0.1908 | 0.3529 |
| 8 | 0.1928 | 0.4806 | 0.6734 | 0.1733 | 0.1532 | 0.3265 |
| 10 | 0.1883 | 0.5030 | 0.6913 | 0.1541 | 0.1545 | 0.3086 |
| 12 | 0.1863 | 0.5219 | 0.7082 | 0.1631 | 0.1287 | 0.2918 |
| 14 | 0.1862 | 0.5327 | 0.7189 | 0.1669 | 0.1142 | 0.2811 |
| 16 | 0.1824 | 0.5486 | 0.731 | 0.1654 | 0.1035 | 0.2689 |

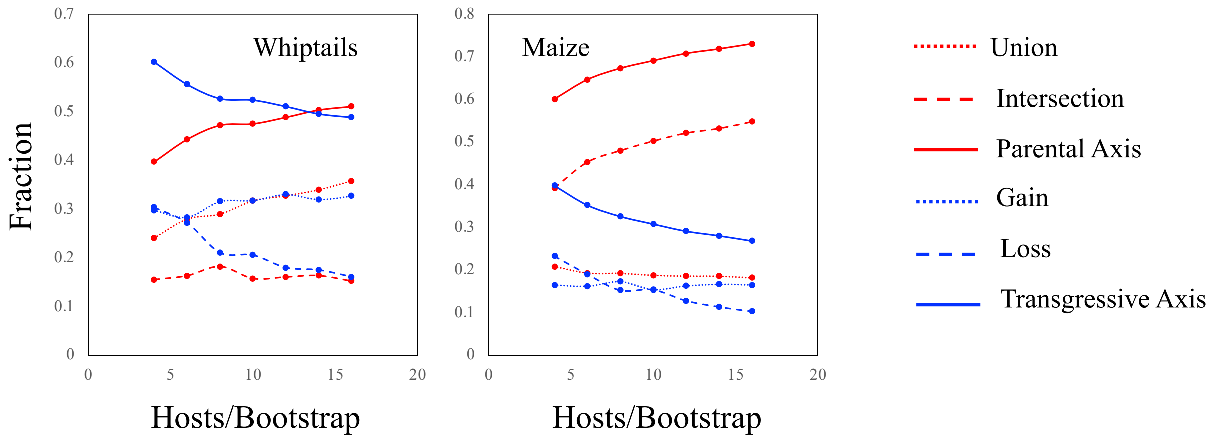

**Fig. S.3.2.** Variation in the 4H index as a function of the number of hosts selected for each bootstrap sample (See Table S3).

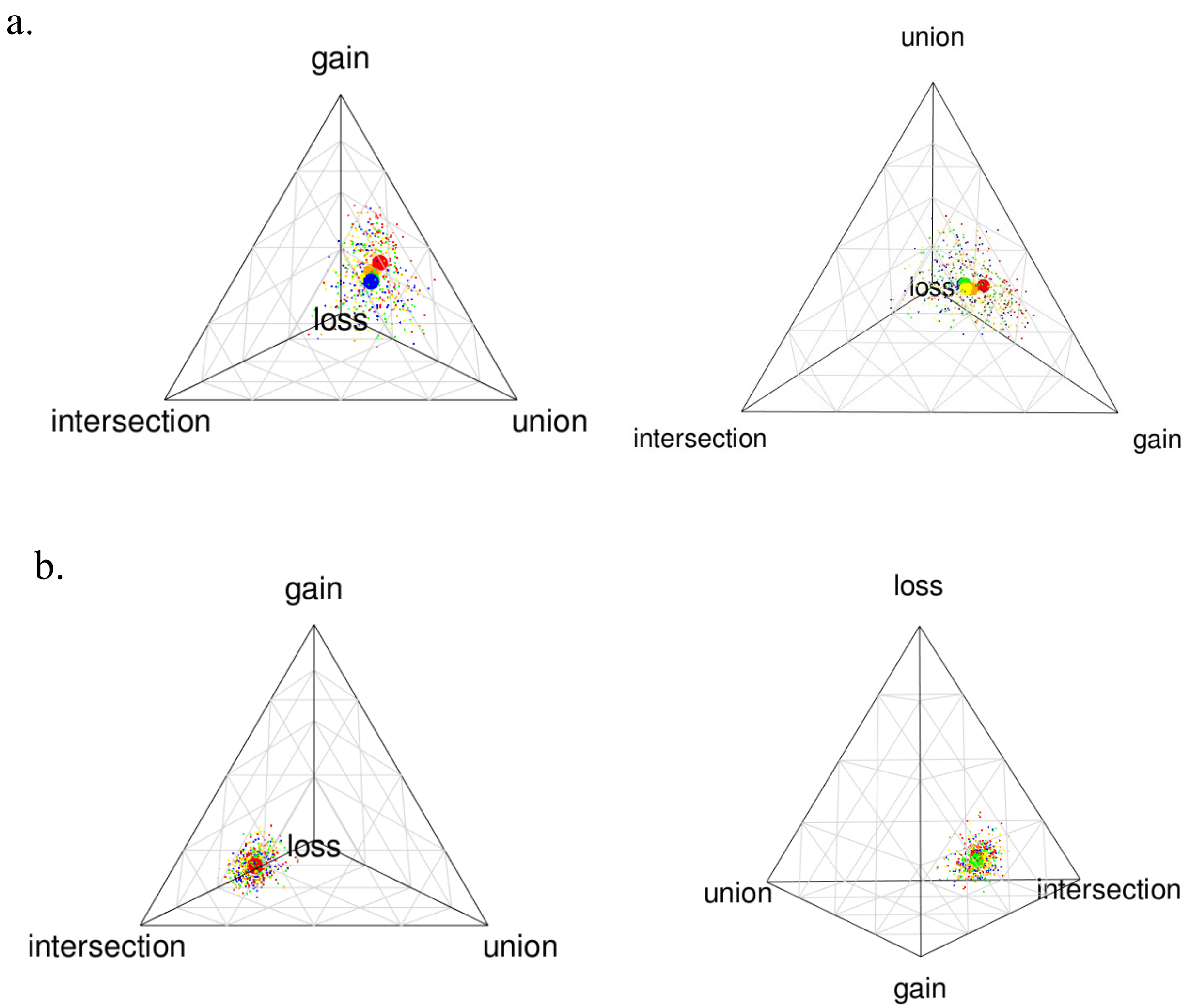

**Fig. S4.1** Quaternary plot showing 100 genus-level bootstrapped samples (small circles) and the bootstrap centroids (large circles) of the 4H index for (a) gut microbiomes from *Aspidoscelis neomexicanus* lizards and (b) B73 line × Mo17 line maize crosses. For the *A. neomexicanus* system, we show samples rarefied to a read depth of 1000 (red), 2000 (orange), 5000 (yellow), 7500 (green) and 10000 (blue). For the maize system, which had a larger number of samples with lower reads, we show samples rarefied to a read depth of 1000 (red), 2000 (orange), 3000 (yellow), 4000 (green) and 5000 (blue). Values of the corresponding centroids and total sums for the parental and transgressive axes are shown, below, in Table S.4. For both systems, we used $\rho = 0.5$, and subsampled 12 individuals of each progenitor and 12 hybrids.

**Table S.4** Variation in the 4H index as a function of read depth. Values are based on 100 genus-level bootstrapped samples of 12 individuals of each progenitor and 12 individuals of the hybrid assuming $\rho = 0.5$ and using the `FourHbootstrap` and `FourHcentroid` functions from the `HybridMicrobiomes` package in R.

| | Parental Axis | | | Transgressive Axis | | |
|---|---|---|---|---|---|---|
| *Aspidoscelis* lizards (16S rRNA gene, cloacal swabs) | | | | | | |
| reads | $\mathcal{U}$ | $\mathcal{I}$ | $\mathcal{U}+\mathcal{I}$ | $\mathcal{G}$ | $\mathcal{L}$ | $\mathcal{G}+\mathcal{L}$ |
| 1000 | 0.3158 | 0.1025 | 0.4183 | 0.3872 | 0.1945 | 0.5817 |
| 2000 | 0.3230 | 0.1303 | 0.4533 | 0.3667 | 0.1801 | 0.5468 |
| 5000 | 0.3148 | 0.1633 | 0.4781 | 0.3506 | 0.1713 | 0.5219 |
| 7500 | 0.3266 | 0.1559 | 0.4825 | 0.3362 | 0.1812 | 0.5174 |
| 10000 | 0.3249 | 0.1598 | 0.4847 | 0.3316 | 0.1837 | 0.5153 |
| B73 line × Mo17 line maize (16S rRNA gene; rhizosphere) | | | | | | |
| reads | $\mathcal{U}$ | $\mathcal{I}$ | $\mathcal{U}+\mathcal{I}$ | $\mathcal{G}$ | $\mathcal{L}$ | $\mathcal{G}+\mathcal{L}$ |
| 1000 | 0.1850 | 0.5142 | 0.6992 | 0.1607 | 0.1402 | 0.3009 |
| 2000 | 0.1849 | 0.5171 | 0.702 | 0.1661 | 0.1319 | 0.298 |
| 3000 | 0.1810 | 0.5225 | 0.7035 | 0.1704 | 0.1260 | 0.2964 |
| 4000 | 0.1872 | 0.5142 | 0.7014 | 0.1702 | 0.1283 | 0.2985 |
| 5000 | 0.1905 | 0.5214 | 0.7119 | 0.1627 | 0.1254 | 0.2881 |

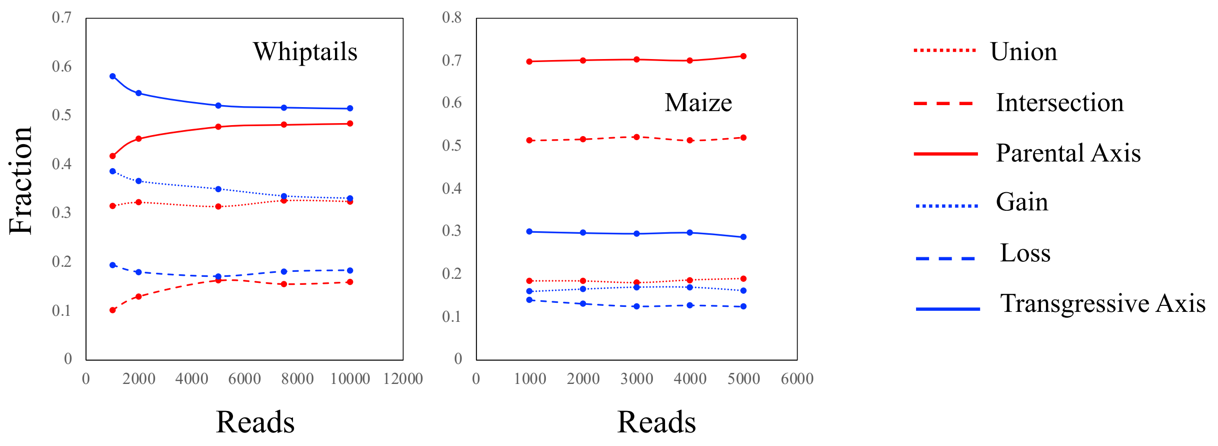

**Fig. S.4.2.** Variation in the 4H index as a function of read depth of the microbiome samples (See Table S.4.).
